# Understanding Influenza A Virus particles detaching from reconstructed cell surfaces

**DOI:** 10.1101/2025.08.06.668852

**Authors:** Thomas Kolbe, Pierre Gaspard, Bortolo Matteo Mognetti

**Affiliations:** Interdisciplinary Center for Nonlinear Phenomena and Complex Systems, Université Libre de Bruxelles (ULB), B-1050 Brussels, Belgium

## Abstract

Influenza infection is a multistage process that involves the trafficking of viral particles across the cell membrane. Before endocytosis, virions target the membrane by binding hemagglutinin ligands to sialic acid residues on cell receptors. After budding, neuraminidase cleaves these residues, enabling virions to detach from the infected cell surface. In this paper, we examine detachment dynamics through simulations and theoretical analysis. We explain experimental findings showing that the time required for virions to detach can decrease as the single-trajectory average number of bridges increases-a counterintuitive result specific to neuraminidase activity. Furthermore, we demonstrate that the detachment time is not governed by a Poisson distribution but depends on multiple factors, including ligand-receptor reaction rates, virion size, and receptor diffusion constant. These results clarify how biochemical parameters regulate the residence time of virions at the cell surface.

**TOC Graphic:** 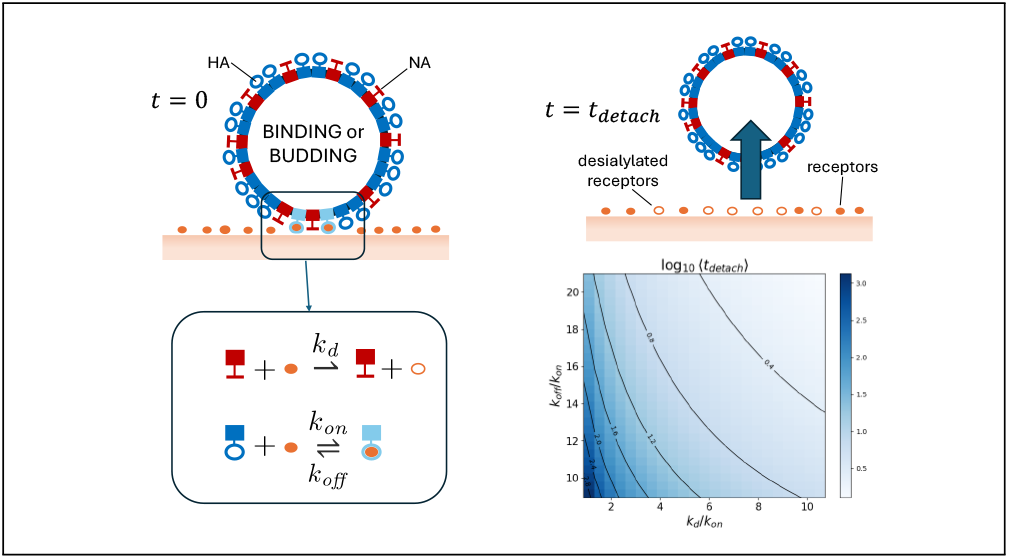

---

Influenza A virus (IAV) is a contagious pathogen with the potential to cause seasonal epidemics and global pandemics. Influenza A (IA) virions (or IAV particles) target the epithelial cells of the respiratory tract. ^1–3^ The dynamics of IA virions in the extracellular medium is regulated by two ligands: hemagglutinin (HA) and neuraminidase (NA). HA ligands bind to sialic acid (SA) residues displayed on cell surface receptors (e.g. glycolipids) and on extracellular molecules such as mucins, which are the main constituents of the mucus. ^4^ While binding to SA residues, HA ligands anchor IA virions to the cell surface through the formation of reversible ligand-receptor bridges (Fig. 1a, d). Instead, NA ligands are enzymes that cleave SA residues from their scaffold. Consequently, NA is an antagonist ligand which limits the number of HA-SA bridges (Fig. 1a, d).

**Figure 1:**
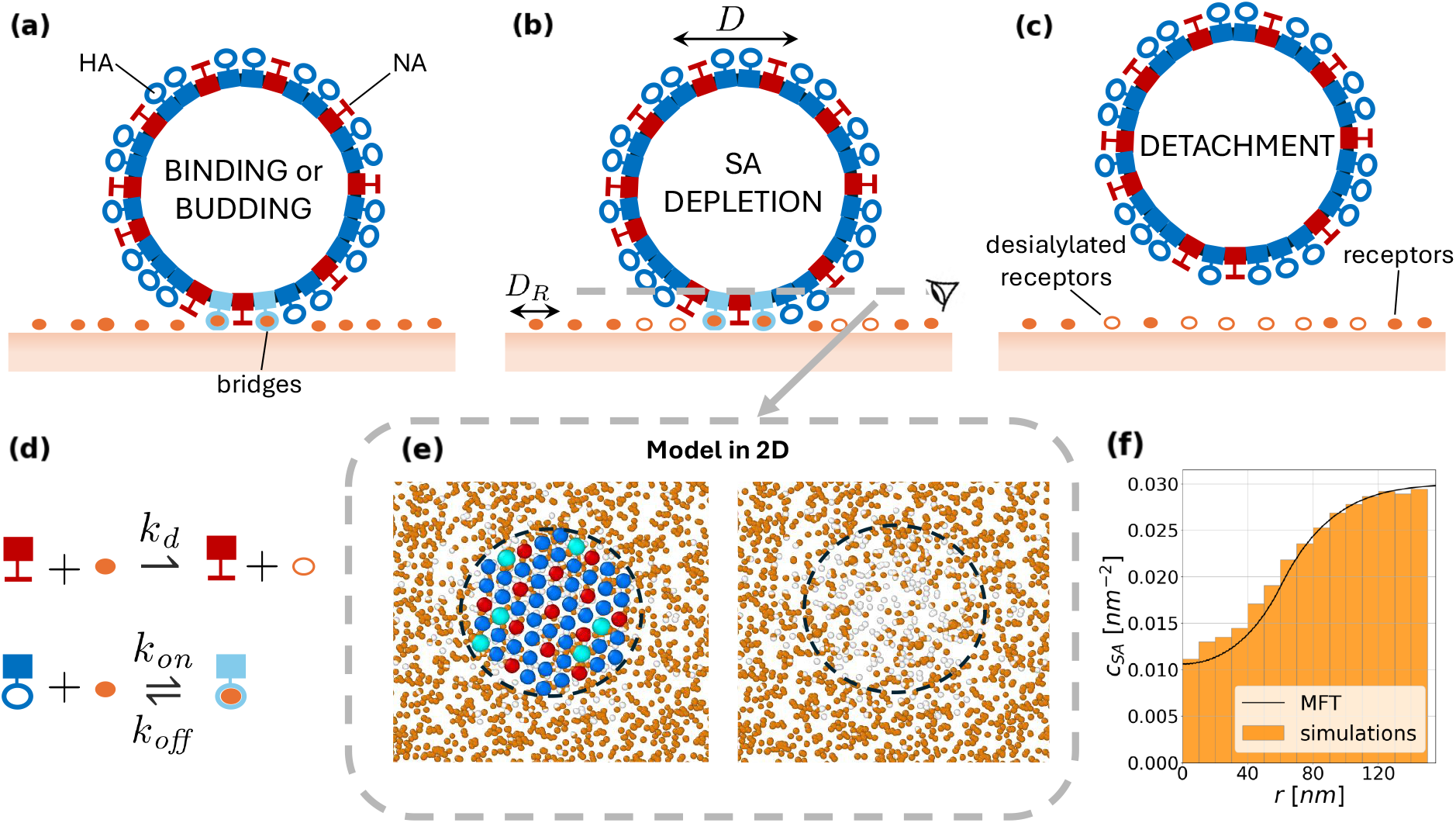
(*a*) Influenza A virions carry NA (red) and HA (blue) ligands binding and cleaving SA molecules (orange dots), tipping the cell’s receptors. (*b*) Virions deplete the local SA concentration, leading to a reduction in the number of bridges and, eventually, to particle detachment (*c*). (*f*) SA concentration as a function of the distance from the center of the particle obtained using simulations and theory for *k*_*d*_ = 3.571 *k*_*on*_, *k*_*off*_ = 16.07 *k*_*on*_ at a time equal to *t* = 2.016*/k*_*on*_. (*d*) Desialylation is controlled by the rate *k*_*d*_ while HA-SA binding and unbinding by *k*_*on*_ and *k*_*off*_, respectively. (*e*) Owing to the slow rotational diffusion constant of the particle, the latter is mapped into a 2D disc that carries receptors found in the particle-surface contact region.

Because of the action of NA enzymes, IAV is an out-of-equilibrium system that constantly consumes chemical fuel (SA residues). Specifically, IAV particles can travel micrometer distances across surfaces functionalized by fixed receptors, ^5–8^ in stark contrast with passive multivalent particles, ^9,10^ which undergo nanometric diffusion when anchored to surfaces by many ligand-receptor bridges.^11–13^ Although the extent to which NA activity facilitates the penetration of the mucus barrier remains under debate, ^7,14,15^ it is generally accepted that coordinated action of NA and HA sustains infection propagation by enhancing virions’ motility.^3,16^ In this manuscript, we study the detachment of virions from cell surfaces. Particle detachment is expected to play an important role in regulating the dynamics of infection spread. ^17,18^

A recent publication by Müller et al. ^19^ reported on how the time taken by a virion to detach from reconstructed cell surfaces (*t*_*detach*_) can increase with the average number of bridges (⟨*n*_*b*_⟩), the latter measured at the single-trajectory level. ^20^ This is a counterintuitive result since, for passive multivalent particles, the number of bridges is correlated with the adhesion strength between the virion and the cell surface. ^9,10^

Here, we use numerical simulations and theory to study *t*_*detach*_. In agreement with Ref.,^19^we prove that the non-monotonicity of *t*_*detach*_ in ⟨*n*_*b*_⟩ is a peculiar non-equilibrium effect of the catalytic activity of NA. This behavior arises from a nonmonotonic trend of the number of HA-SA bridges (*n*_*b*_) that features a local maximum when plotted against time. In general, *t*_*detach*_ is a stochastic variable that does not follow a Poisson distribution but is controlled by multiple timescales, including the diffusion constant of the receptors and ligands’ reaction rates. Our results provide a biophysical explanation of how the synergistic activity between NA and HA ligands could be pivotal for infection spread.

## The model and the simulation algorithm

Virions are mapped into rigid particles while the ligands/receptors into points decorating the particle’s surface/surface. Ligands can only interact with receptors found at a distance smaller than *λ* (*λ* = 7.5 nm^7,8^). Given that the interaction range *λ* is much smaller than the virion’s radius, only ligands in a patch facing the surface can interact with the receptors.

Therefore, for simplicity, we employ the model of Ref. ^8^ (Fig. 1e) in which the interacting patch of the virion is mapped into a 2D disc of radius *R* (*R* = 60 nm unless specified). This 2D model is reasonable given that typical detachment times *t*_*detach*_ ^19^ are much smaller than the timescales at which the virions are expected to rotate.^1^ The disc carries *N*_*HA*_ hemagglutinin (*N*_*HA*_ = 48) and *N*_*NA*_ neuraminidase ligands (*N*_*NA*_ = 13) arranged as in a triangular lattice (Fig. 1e). NA ligands are uniformly scattered over the available lattice sites. couple of ligand-receptor molecules found at a lateral distance smaller than *λ* (*λ* = 7.5 nm) can react (Fig. 1*a*): HA can bind or unbind SA molecules with a rate constant equal to, respectively, *k*_*on*_ and *k*_*off*_ (Fig. 1*d*). Instead, NA ligands can cleave SA molecules from the receptors at a rate constant equal to *k*_*d*_ (Fig. 1*d*). Receptors are uniformly distributed over the surface with a starting density equal to 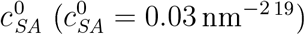 and move with a diffusion constant equal to *D*_*R*_ (*D*_*R*_ = 0.3 *μ*m^2^/s unless specified, Fig. 1b). Due to the reversibility of bridges and the action of the NA, the number of bridges (*n*_*b*_) is affected by stochastic fluctuations that eventually lead to the detachment of the particle from the surface when the number of bridges becomes equal to zero, *n*_*b*_ = 0.

We employ a reaction-diffusion algorithm. At each simulation step, a list of reactions (binding/unbinding or desialylation events) is implemented according to Gillespie’s algorithm. ^22^ Receptors and the disc are diffused using Brownian dynamics. With the particle, we also diffuse the bound receptors. For the particle, we rely on the observation of Ref.^20^ and employ a diffusion constant inversely proportional to the number of bridges featured by the particle at a given time, *D* = *D*_*R*_*/n*_*b*_. This expression shows how the friction exerted by the lipid bilayer onto the particle is additive in the number of bridges, in contrast with Saffman and Delbrück theory. ^232^ Importantly, as discussed below, our results are not impacted by the specific form of the diffusion constant *D* employed (SI Fig. 3). Further details about our simulation protocol and its robustness are reported in SI Sec. 1. ^3^

## Simulation results

Fig. 2 reports the predictions of the detachment time *t*_*detach*_ for three different values of the desialylation rate *k*_*d*_. Each data point corresponds to a single simulation while lines are the results of the theory developed below. The variability of *t*_*detach*_ obtained with a set of simulations employing the same system parameters highlights the stochastic nature of *t*_*detach*_. Fig. 2a correlates *t*_*detach*_ with the single-trajectory average of the number of bridges ⟨*n*_*b*_ ⟩. In agreement with experiments ^19^ we find that *t*_*detach*_ decreases with ⟨*n*_*b*_⟩. The trend arises from the activity of the NA ligands and disappears when *k*_*d*_ = 0 (SI Fig. 4). Fig. 2b reports the detachment time distributions obtained from the three sets of data of Fig. 2a. These distributions deviate from a Poissonian function (as found for *k*_*d*_ = 0 SI Fig. 5). In particular they feature a global maximum.

**Figure 2:**
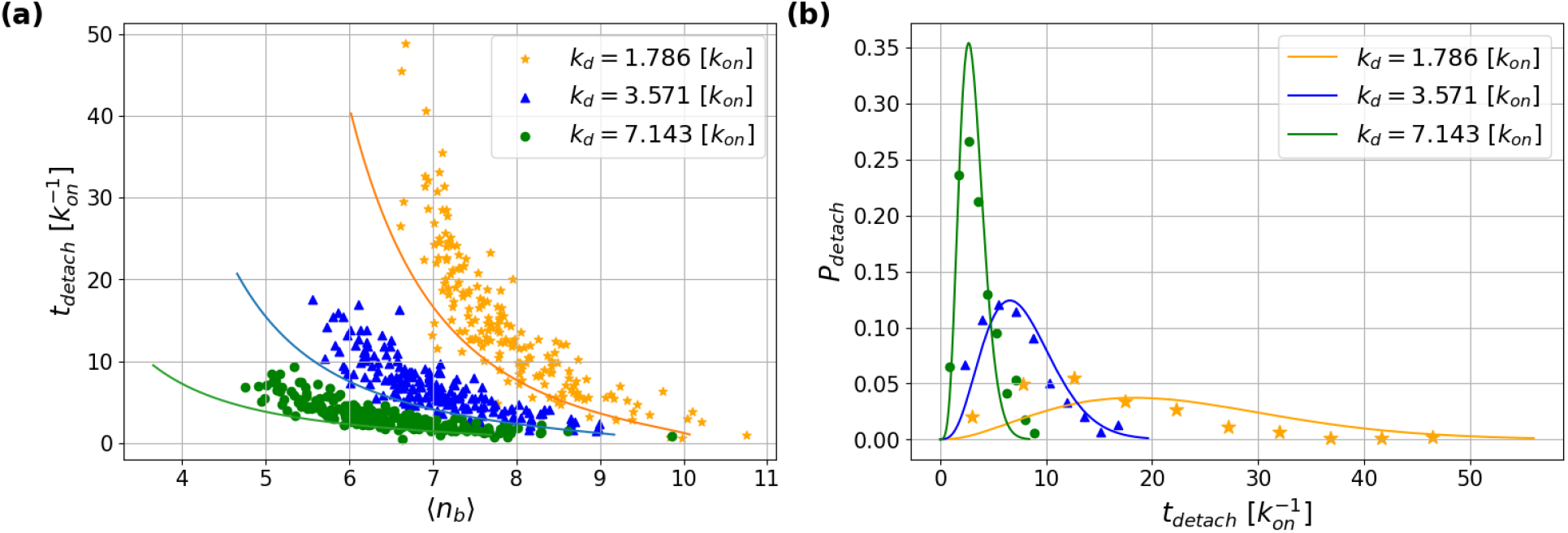
(*a*) Detachment times *versus* single-trajectory average number of bridges ⟨*n*_*b*_ ⟩ for three different desialylation rates. For each value of *k*_*d*_ we run 200 simulation repeats. (*b*) Distributions of the detachment times corresponding to the three data sets considered in panel (*a*). We used *k*_*off*_ = 16.07 *k*_*on*_.

To rationalize the finding of Fig. 2 in Fig. 3 we study the number of bridges *n*_*b*_ versus time using ten simulations with *k*_*d*_ = 3.571 *· k*_*on*_ (colored lines). Starting from a configuration with *n*_*b*_ = 0, the number of bridges readily peaks to a global maximum before beginning to decrease. At the single trajectory level, while decreasing, *n*_*b*_ fluctuates rapidly around a baseline that changes on longer timescales. The fluctuations around the baseline underlie the stochastic nature of *t*_*detach*_. In Fig. 3, we model the baseline using the predictions of the theory developed below. The rapid fluctuations of the simulation trajectories are due to HA-SA binding/unbinding events regulated by *k*_*on*_ and *k*_*off*_. Instead, the slow decrease of the baseline is due to the depletion of SA ligands available to form bridges and is controlled by *k*_*d*_ and *D*_*R*_, as discussed below. The local maximum in *n*_*b*_ is peculiar to the presence of activity in the system SI Fig. 6). The local maximum of Fig. 3 rationalises the decrease of the detachment time with ⟨ *n*_*b*_ ⟩ reported in Fig. a. In particular, ⟨*∗n*_*b*_*⟩* decreases with *t*_*detach*_ when *t*_*detach*_ is greater than the value that maximises *n*_*b*_.

**Figure 3:**
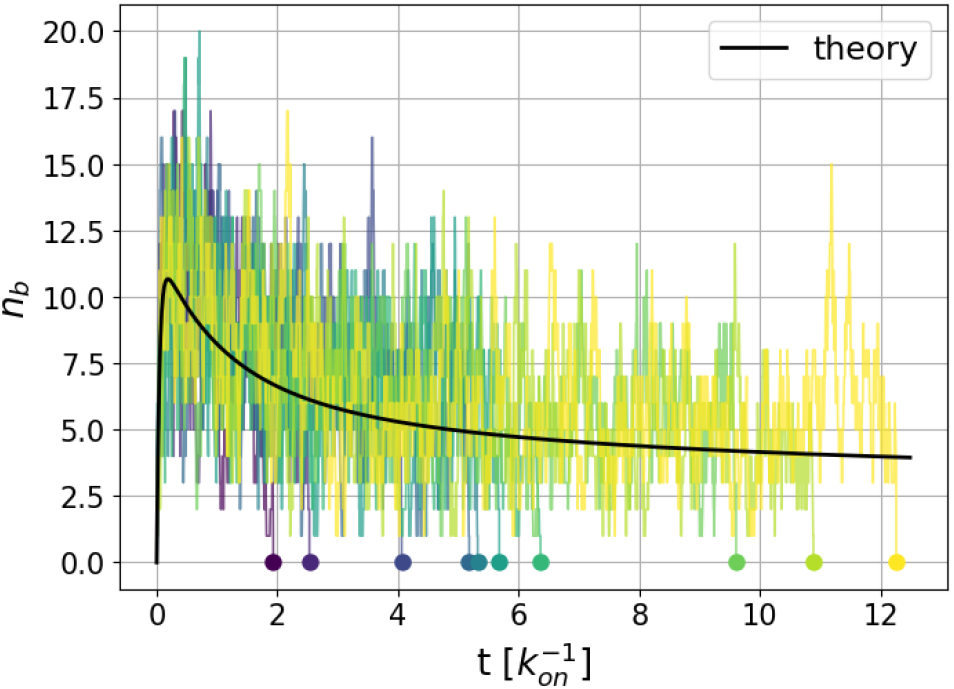
Number of bridges *versus* time. Colored plots refer to simulation trajectories, while the black line refers to the theory’s prediction. Dots on the temporal axis correspond to simulation detachment times. We used *k*_*d*_ = 3.571 *k*_*on*_ and *k*_*off*_ = 16.07 *k*_*on*_.

## Mean field theory

To corroborate the previous simulation results, we develop a mean-field theory to calculate the evolution of the number of bridges baseline in Fig. 3) and the distribution of the detachment time (Fig. 2b).^4^ We distribute HA and NA ligands uniformly over the disc with coverage equal to

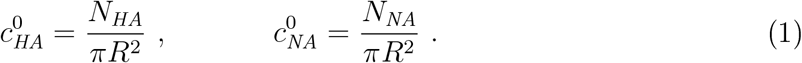

Similarly, we define by *c*_*SA*_(**x**, *t*) and *c*_*B*_ (**x**, *t*), respectively, the concentration of free SA (i.e, not bound to HA) and bridges on the surface with an initial value equal to 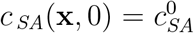 and *c*_*B*_ (**x**, 0) = 0 The evolution of *c*_*SA*_ and *c*_*B*_ is then given by the following coupled partial differential equations:

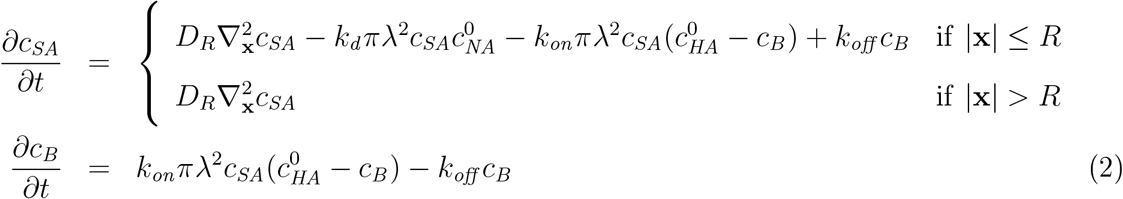

where 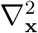 is the 2D Laplacian and *c*_*B*_ (**x**, *t*) = 0 if |**x**| *> R* or *t* = 0 The center of the disc is fixed at **x** = (0, 0), and we assume cylindrically symmetric solutions for *c*_*SA*_ and *c*_*B*_ (this assumption is discussed below). Eqs. 2 are solved numerically within a disk of radius *L* (*L* ≫ *R*) using Neumann boundary conditions (the analysis of the solutions with Dirichlet boundary consitions is presented in SI Sec. 3). Neumann boundary conditions set to zero the inflow current of SA receptors from the boundary of the system. This is consistent with *in vitro* experiments without SA replenishment.

The theory is in quantitative agreement with simulations, as shown in Fig. 1f for a snapshot of the SA concentration, *c*_*SA*_, and in Fig. 3 for the baseline of the number of bridges, *n*_*b*_(*t*), the latter calculated by integrating the bridge s distribution, *n*_*b*_(*t*) = ∫*d***x** *c*_*B*_(**x**, *t*). By analysing the asymptotic solutions of Eqs. 2 (see SI Sec. 2), we can prove that, after reaching the peak, the number of bridges follows a multi-regime decay characterized by two exponential functions, *n*_*b*_(*t*) ≈ exp(−*t*/*τ*_*i*_) with *i* = 1, 2 and

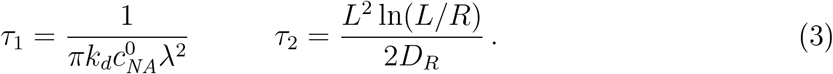

Immediately after the peak, *n*_*b*_ decays with a timescale equal to *τ*_1_, which is only controlled by the desilylation rate *k*_*d*_ (Fig. 4b). At later stages of the dynamics, the decay is controlled by *τ*_2_, which depends on the diffusion constant *D*_*R*_, the radius of the particle *R*, and the size of the system, *L* (Fig. 4a). Figs. 4 show how Eqs. 3 are semi-quantitative, especially for the first decay, which is, in fact, affected by SA replenishment (see SI Sec. 2 and SI Fig. 2). Importantly, while the size of the particle does not affect the early-stage decay rate, it does affect the height of the peak. Accordingly, the detachment time *t*_*detach*_ is expected to increase significantly as a function of the particle size *R* (SI Fig. 7). The pleiomorphic nature of IAV is therefore likely to result in a broad spectrum of detachment times.

**Figure 4:**
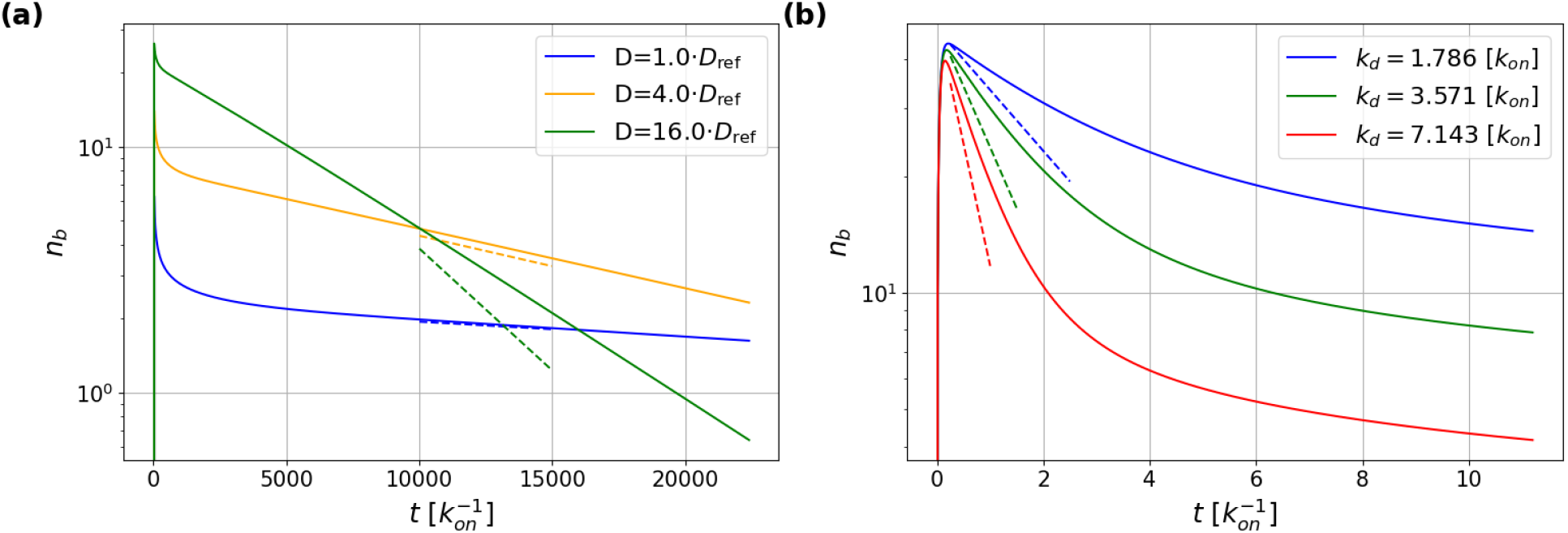
Theoretical predictions (full lines) and scaling analysis (dashed lines) for the late (*a*) and the early decay (*b*) of the number of bridges *verses* time. The scaling analysis is based on Eqs. 3. In (*a*) we used *k*_*d*_ = 3.571 *k*_*on*_ with *D*_ref_ = 0.3 *μm*^2^ In both panels, we used *k*_*off*_ = 16.07 *k*_*on*_ and *R* = 120 nm.

The results of Eq. 3 assume that the equilibration time of the number of bridges at a given concentration of SA is much faster than the timescales controlling the relaxation of *c*_*SA*_ (Fig. 3 and SI Sec. 2.2). Within the same approximation, we can derive an expression for the distribution of the detachment times *P*_*detach*_ (*t*_*detach*_): At a given SA concentration *c*_*SA*_; we can employ first passage theory ^25^ to calculate the mean time, *τ*_*d*_ (*t*); taken for the particle to reach the *n*_*b*_ = 0 condition, *τ*_*d*_ (*t*) = 1*/k*_*det*_ (*t*). *k*_*det*_ (*t*) is the rate at which a passive multivalent particle would detach from a surface carrying a concentration of SA equal to *c*_*SA*_ (**x**, *t*); ^26^ the latter given by Eq. 2. Details about the calculation of *k*_*det*_ are reported in SI Sec. 4. The detachment distribution function is then given by a non-homogeneous Poissonian process

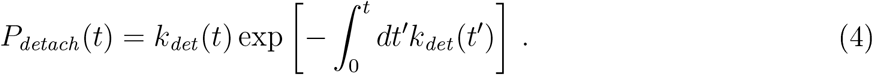

Fig. 2b reports the predictions of Eq. 4 as compared to simulation results. Overall, we find quantitative agreement between simulations and theory. This result validates the proposed theory as a tool to extensively study the detachment time.

### Discussion and conclusions

Our mean field theory corroborates the model presented in Fig. 1a-c in which virions detach from the cell surface as a result of a local depletion in the sialic acid concentration at the cell surface, leading to a decrease in the avidity and therefore in the number of bridges. The unexpected anticorrelation between the detachment time, *t* _*detach*_, and the single-trajectory average number of bridges, ⟨*n*_*b*_⟩,^19^ is a purely statistical effect due to the presence of a global maximum in the number of bridges *n*_*b*_ *versus* time (Fig. 3). This result is peculiar to the presence of the enzymatic activity of the NA ligands, while it is not sensitive to the motility of the virion. The latter statement is proven in SI Fig. 3 where we repeat simulations using a constant diffusion constant *D* for the particle (instead of *D ~ D*_*R*_*/n*_*b*_).

The detachment time is controlled by the HA-NA affinity (*k* _*on*_ */k*_*off*_), the size of the particle *R*, and the desilylation rate *k*_*d*_. *R* and *k*_*on*_ */k*_*off*_ affect the maximum number of HA-SA bridges formed (height of the peak in Fig. 3), while *k*_*d*_ the timescale, *τ*_1_ (Eq. 3), controlling the early decay of *n*_*b*_. The detachment time is not affected by the diffusion of the receptors *D*_*R*_ if a virion lowers the number of bridges enough before reaching the crossover towards the second decay controlled by *τ*_2_. High values of the diffusion constant, *D*_*R*_, tend to increase *n*_*b*_ at short timescales by favouring the replenishment of SA receptors (Fig. 4a). The time at which this replenishment becomes important controls the crossover between the two exponential decays of Eq. 3. At later stages of the dynamics, *D*_*R*_ decreases the number of bridges by decreasing the timescale *τ*_2_ (Eq. 3), controlling the second decay of *n*_*b*_. It should be noted that in the second decay timescale, the SA concentration becomes depleted over the entire cell. Such a configuration becomes more probable in the presence of many virions. The many-virion case can be studied using 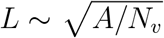 in Eq. 3, if *A* is the area of the cell and *N*_*v*_ the number of attached virions.

In Fig. 5 we use Eq. 4 to calculate the detachment time as a function of the reaction rates. We find that *t*_*detach*_ can vary by orders of magnitude as the rates change within experimentally plausible ranges. ^27–29^ It is believed that the most infectious strains of IAV are those in which the activity of NA and HA ligands is balanced. The present study shows that such a balance, along with other relevant parameters (like *D*_*R*_ and *R*), plays a role in controlling the residence time of a virion on the surface of the cell before detaching. IAV particle residence time is expected to be regulated as it controls the efficiency of endocytosis (in which the virions need to recruit specific receptors to trigger entrance ^30^) and trafficking of virions towards uninfected cells. In the future, it will be important to consider the results of our study, distilled into the presented theory, to study infection spreading at the tissue level.

**Figure 5:**
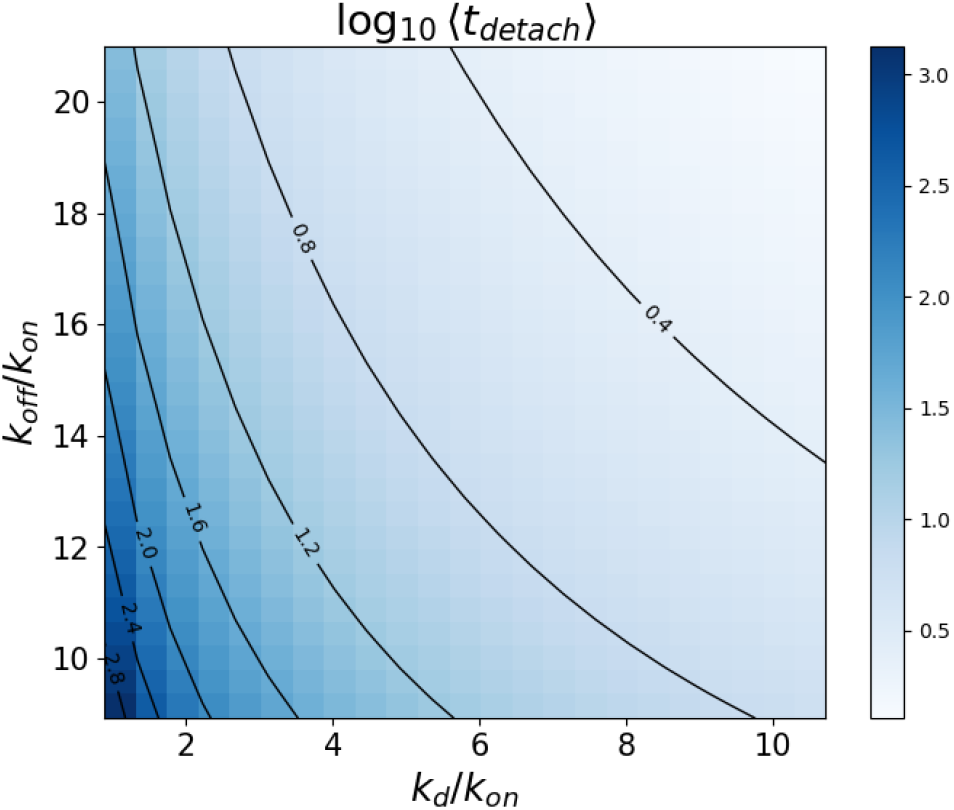
Average value of the detachment time; ⟨*t*_*detach*_⟩ as a function of the reaction rates. The average has been obtained using Eq. 4 as ⟨*t*_*detach*_⟩ = ⎰*dt P*_*detach*_ (*t*) *· t*.

## Acknowledgenent

B.M.M. and T.K. are supported by a PDR grant of the FRS-FNRS (Grant No. T021024F). Computational resources have been provided by the Consortium des Équipements de Calcul Intensif (CECI), funded by the Fonds de la Recherche Scientifique de Belgique (F.R.S.-FNRS) under Grant No. 2.5020.11 and by the Walloon Region.

## Supporting Infornation Available

The following files are available free of charge.

- *SI-achemso. Pdf*: This file contains supplementary figures. It also provides details about the simulation algorithm and the theoretical developments leading to Eqs. 3 and Eq. 4.

## Footnotes

1 The latter is expected to be controlled by the diffusion constants of particles moving across surfaces with fixed receptors ^21^ which are different orders of magnitude smaller than the diffusion constants observed in Refs. ^19,20^

2 Note that Saffman and Delbruck theory has been proven more adequate than an additive friction term when studying the dynamics of micron-sized particles. ^24^

3 The simulation program is available at the following URL: https://github.com/tkol98/IAV_detachment

4 The scripts implementing the theory are available at the following URL: https://github.com/bmognetti/IAV_detachment_dynamics.

## References

(1) Bouvier, N. M.; Palese, P. The biology of influenza viruses. 26, D49–D53.

(2) Dou, D.; Revol, R.; Östbye, H.; Wang, H.; Daniels, R. Influenza A Virus Cell Entry, Replication, Virion Assembly and Movement. 9, Publisherf Frontiers.

(3) de Vries, E.; Du, W.; Guo, H.; de Haan, C. A. Influenza A virus hemagglu-tinin-neuraminidase-receptor balance: Preserving virus motility. 28, 57–67, Publisher: Elsevier.

(4) Wallace, L. E.; Liu, M.; van Kuppeveld, F. J. M.; de Vries, E.; de Haan, C. A. M. Respiratory mucus as a virus-host range determinant. 29, 983–992.

(5) Sakai, T.; Nishimura, S. I.; Naito, T.; Saito, M. Influensa A virus hemagglutinin and neuraminidase act as novel motile machinery. 7, Publisher: Nature Publishing Group.

(6) Guo, H.; Rabouw, H.; Slomp, A.; Dai, M.; van der Vegt, F.; van Lent, J. W. M.; McBride, R.; Paulson, J. C.; de Groot, R. J.; van Kuppeveld, F. J. M.; de Vries, E.; de Haan, C. A. M. Kinetic analysis of the influensa A virus HA/NA balance reveals contribution of NA to virus-receptor binding and NA-dependent rolling on receptor-containing surfaces. 14, 1–31, Publisher: Public Library of Science.

(7) Vahey, M.; Fletcher, D. A. Influensa A virus surface proteins are organised to help penetrate host mucus. 8, e43764, Publisher: eLife Sciences Publications Limited.

(8) Stevens, L.; Buyl, S. d.; Mognetti, B. M. M. The sliding motility of the bacilliform virions of lnfluensa A viruses. 19, 4491–4501, Publisher: The Royal Society of Chemistry.

(9) Bell, G. I. Models for the specific adhesion of cells to cells: a theoretical frame ork for adhesion mediated by reversible bonds between cell surface molecules. Science 1978, 200, 618–627.

(10) Martines-Veracoechea, F. J.; Frenkel, D. Designing super selectivity in multivalent nano-particle binding. 108, 10963–10968, Publisher: Proceedings of the National Academy of Sciences.

(11) Martines-Veracoechea, F. J.; Mognetti, B. M.; Angioletti-Uberti, S.; Varilly, P.; Frenkel, D.; Dobnikar, J. Designing stimulus-sensitive colloidal walkers. 10, 3463–3470, Publisher: Royal Society of Chemistry.

(12) Marbach, S.; Zheng, J. A.; Holmes-Cerfon, M. The nanocaterpillar’s random walk: diffusion with ligand-receptor contacts. 18, 3130–3146, Publisher: Royal Society of Chemistry.

(13) Lowensohn, J.; Stevens, L.; Goldstein, D.; Mognetti, B. M. Sliding across a surface: Particles with fixed and mobile ligands. The Journal of Chemical Physics 2022, 156.

(14) Fang Steukers, L.; Forier Xiong, R.; Braeckmans K.; Reeth, K. V.; Nauwynck, H. A Beneficiary Role for Neuraminidase in Influenza Virus Penetration through the Respiratory Mucus. 9, e110026, Publisher: Public Library of Science.

(15) aler, L.; Iverson, E.; Bader, S.; Song, D.; Scull, M. A.; Duncan, G. A. Influenza A virus diffusion through mucus gel networks. 5, 1–9, Publisher: Nature Publishing Group.

(16) Exploring virus release as a bottleneck for the spread of influenza A virus infection in vitro and the implications for antiviral therapy with neuraminidase inhibitors. 12.

(17) Jacobs, N. T.; Onuoha, N. O.; Antia, A.; Steel, J.; Antia, R.; Lowen, A. C. Incomplete influenza A virus genomes occur frequently but are readily complemented during localized viral spread. 10, 3526, Publisher: Nature Publishing Group.

(18) Rudiger, D.; upke, S.; Laske, T.; Zmora, P.; Reichl, U. Multiscale modeling of influenza A virus replication in cell cultures predicts infection dynamics for highly different infection conditions. 15, e1006819, Publisher: Public Library of Science.

(19) Muller, M.; Lauster, D.; Wildenauer, H. H.; Herrmann, A.; Block, S. Mobility-based quantification of multivalent virus-receptor interactions: new insights into influenza A virus binding mode. Nano letters 2019, 19, 1875–1882.

(20) Block, S.; Zhdanov, V. P.; Hook, F. Quantification of multivalent interactions by tracking single biological nanoparticle mobility on a lipid membrane. Nano letters 2016, 16, 4382–4390.

(21) Jana, P. K.; Mognetti, B. M. Translational and rotational dynamics of colloidal particles interacting through reacting linkers. Physical Review E 2019, 100, 060601.

(22) Gillespie,. T. Exact Stochastic Simulation of Coupled Chemical Reactions. 81, 2340–2361.

(23) Saffman, P.; Delbrück, M. Brownian motion in biological membranes. Proceedings of the National Academy Sciences 1975, 72, 3111–3113.

(24) Merminod, S.; Edison, J. R.; fang, H.; Hagan, M. f.; Rogers, W. B. Avidity and surface mobility in multivalent ligand-receptor binding. Nanoscale 2021, 13, 12602–12612.

(25) Van kampen, N. G. In Stochastic processes in Physics and chemistry (Third Edition); Van kampen, N. G., Ed.; North-Holland Personal Library; Elsevier, pp 292–325.

(26) Erdmann, T. S.; Schwarz, U. S. Impact of receptor-ligand distance on adhesion cluster stability. 22, 123–137.

(27) Takemoto,. K.; Skehel, J. J.; Wiley, D. C. A Surface Plasmon Resonance Assay for the Binding of nfiuenna Virus Hemagglutinin to Its Sialic Acid Receptor. 217, 452–458.

(28) Benton,. J.; Martin, S. R.; Wharton, S. A.; McCauley, J. W. Biophysical Measurement of the Balance of Influenza A Hemagglutinin and Neuraminidase Activities. 290, 6516–6521.

(29) Liu, S.-L.; Zhang, Z.-L.; Tian, Z.-Q.; Zhao, H.-S.; Liu, H.; Sun, E.-Z.; Xiao, G. F.; Zhang, W.; Wang, H.-Z.; Pang,. D.-W. Effectively and Effciently Dissecting the Infection of Influenza Virus by Quantum-Dot-Based Single-Particle Tracking. 6, 141–150, Publisher: American Chemical Society.

(30) Broich, L.; Wullenkord, H.; Osman, M. K.; Fu, Y.; Müsken, M.; Reuther, P.; Brönstrup, M.; Sieben, C. Single infiuenna A viruses induce nanoscale cellular reprogramming at the virus-cell interface. 16, 3846, Publisher: Nature Publishing Group.

