## Supporting Information for "Understanding Influenza A Virus particles detaching from reconstructed cell surfaces"

#### 1 Simulation protocol

When sampling the detachment time, $t_{detach}$ (see Main Fig. 2), we initialize the particle in an unbound state (with $n_b = 0$) at $t = 0\,\mathrm{s}$. At this stage, the disc diffuses freely with $D = D_R$ until the first bridge is formed. The simulation continues until the number of bridges returns to zero (for $t = t_{detach}$), at which point the particle is considered detached, and the simulation ends. In the analysis of Main Fig. 2, we discard trajectories that detach immediately after binding (with $\langle n_b \rangle < 4$). This is consistent with experimental analysis[1] in which virions are considered bound only when detected on the surface for a sufficiently long time ($t_{detach} > 0.5\,\mathrm{s}$[1]).

To probe the robustness of our simulation algorithm, in SI Fig. S1 we demonstrate that the calculation of $t_{detach}$ is not affected by the algorithm's time step $\Delta t$.

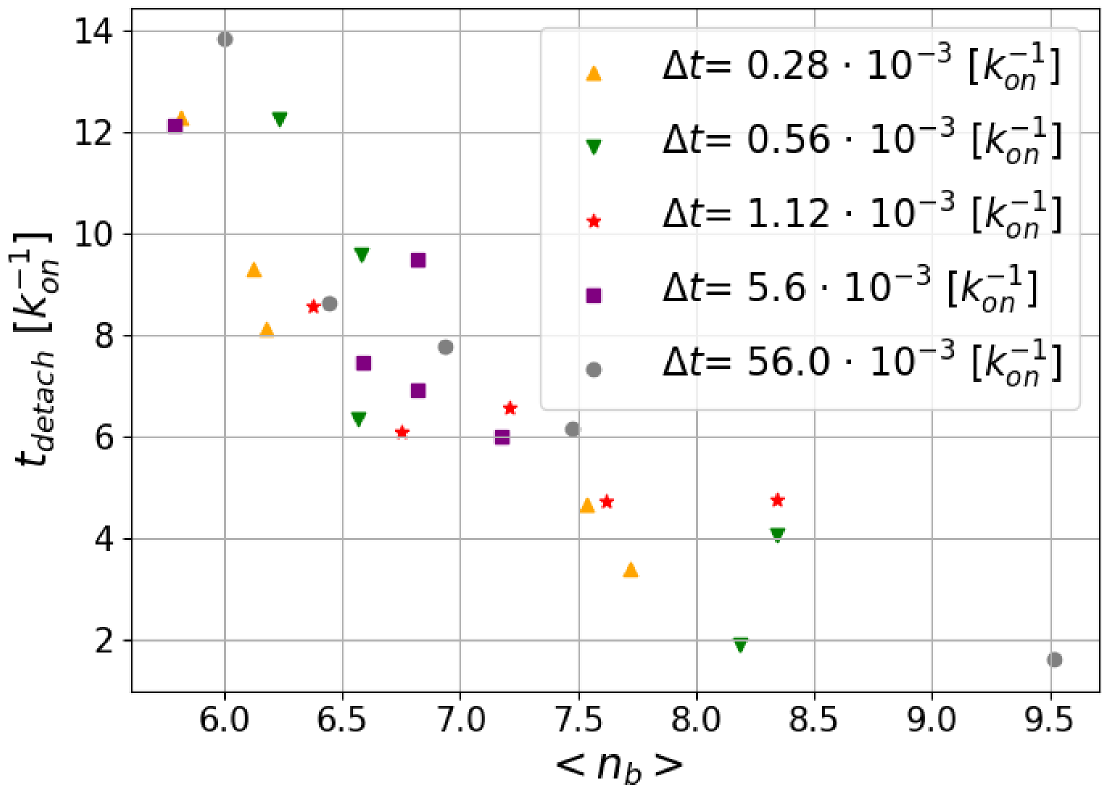


Figure S1: Detachment time *vs* $\langle n_b \rangle$ for different time steps $\Delta t$. Simulation parameters: $k_d = 3.571\, k_{on}$, $k_{off} = 16.07\, k_{on}$.

#### 2 Asymptotic analysis of the mean-field theory: Neumann boundary conditions

Main Eqs. 2 read as

$$\partial_t\, c_{SA} = D\nabla^2 c_{SA} - K_d\, c_{SA} - K_{on}\left(1 - \frac{c_B}{c^0_{HA}}\right) c_{SA} + k_{off}\, c_B$$
$$\partial_t\, c_B = K_{on}\left(1 - \frac{c_B}{c^0_{HA}}\right) c_{SA} - k_{off}\, c_B \qquad (1)$$

for $r < R$, and

$$\partial_t c_{SA} = D\nabla^2 c_{SA} \tag{2}$$

$$c_B = 0 \tag{3}$$

for $R < r < L$. At $r = R$, the function $c_{SA}$ and its derivative $\partial_r c_{SA}$ should be continuous.

The equations should be solved in two dimensions, within a disk of radius equal to $L$ ($L > R$), with the zero-flux boundary condition (Neumann boundary conditions)

$$\partial_r c_{SA}(r = L, t) = 0\,. \tag{4}$$

The parameters of the PDEs are defined as

$$\begin{aligned} K_d &\equiv k_d \pi \lambda^2 c^0_{NA} = k_d \left(\frac{\lambda}{R}\right)^2 N_{NA} \\ K_{on} &\equiv k_{on} \pi \lambda^2 c^0_{HA} = k_{on} \left(\frac{\lambda}{R}\right)^2 N_{HA} \end{aligned} \tag{5}$$

where $D$ is the diffusion constant of the receptors in the reference frame of the particle. For simplicity, we set $D = D_R$.

#### 2.1 The initial conditions

The initial conditions are fixed to

$$c_{SA}(\mathbf{r}, t = 0) = c^0_{SA} \qquad \text{and} \qquad c_B(\mathbf{r}, t = 0) = 0\,. \tag{6}$$

As a consequence, $c_B$ undergoes a rapid increase at early time, according to $\partial_t\, c_B = K_{on} c^0_{SA}$, over a very short time interval. Thereafter, $c_B$ reaches a maximum and slowly decays. This maximum can be used as a new initial condition for the slow decay. The maximum of $c_B$, where $\partial_t c_B = 0$, is given by

$$c_B|_{max} = \frac{K_{on}\left(1+\frac{c^0_{SA}}{c^0_{HA}}\right) + k_{off} - \sqrt{K^2_{on}\left(1-\frac{c^0_{SA}}{c^0_{HA}}\right)^2 + k^2_{off} + 2k_{off}K_{on}\left(1+\frac{c^0_{SA}}{c^0_{HA}}\right)}}{2\frac{k_{on}}{c^0_{HA}}} \tag{7}$$

which in the weak-binding regime becomes

$$c_B|_{max} \simeq \frac{K_{on}}{k_{off}}\, c^0_{SA}\,. \tag{8}$$

The previous expression neglects desialylation (controlled by $k_d$) and the diffusion of the receptors (controlled by $D_R$), replenishing the concentration of free SA. Such replenishment is already at play during the formation of the peak in $n_b$ as shown in Fig. S2.

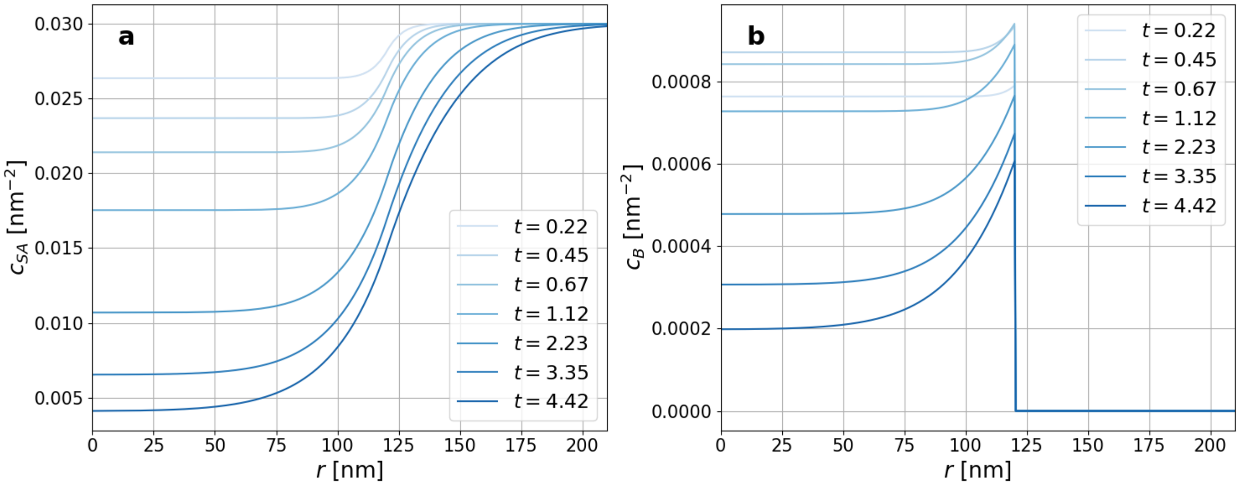


Figure S2: Concentration of SA (a) and bridges (b) at different times (the maximum number of bridges is reached for $t \approx 0.45$). Times are expressed in units of $10^{-6}/k_{on}$. Simulation parameters: $R = 120\,\mathrm{nm}$, $k_{off} = 16.07/k_{on}$, $k_d = 3.57/k_{on}$.

#### 2.2 Linearization around the stationary solution

The stationary solution, such that $\partial_t c_{SA} = 0$ and $\partial_t c_B = 0$, is the uniform solution given by

$$c_{SA} = 0 \qquad \text{and} \qquad c_B = 0\,. \tag{9}$$

Around this solution, the linearized equations read

$$\partial_t \, c_{SA} = D\nabla^2 c_{SA} - (K_d + K_{on}) \, c_{SA} + k_{\mathit{off}} \, c_B \tag{10}$$

$$\partial_t \, c_B = K_{on} \, c_{SA} - k_{\mathit{off}} \, c_B \tag{11}$$

for $r < R$, and

$$\partial_t c_{SA} = D\nabla^2 c_{SA} \tag{12}$$

$$c_B = 0 \tag{13}$$

for $R < r < L$.

The equation for $c_B$ can be solved using only an integration over time, which gives

$$c_B(\mathbf{r}, t) = e^{-k_{\mathit{off}} t} c_B(\mathbf{r}, 0) + K_{on} \int_0^t d\tau \; e^{-k_{\mathit{off}}(t-\tau)} c_{SA}(\mathbf{r}, \tau) \tag{14}$$

for $r < R$, and $c_B = 0$ for $R < r < L$.

Since the typical timescale controlling bridge opening

$$t_{\mathit{off}} \equiv \frac{1}{k_{\mathit{off}}} \tag{15}$$

is the shortest time scale of the problem, we can make the following approximation,

$$c_B(\mathbf{r}, t) \simeq \frac{K_{on}}{k_{\mathit{off}}} \, c_{SA}(\mathbf{r}, t) \tag{16}$$

for $r < R$. Substituting this approximation into the evolution equation for $c_{SA}$, we obtain the following simplified equation,

$$\boxed{\partial_t \, c_{SA} = D\nabla^2 c_{SA} - K_d \, \theta(R - r) \, c_{SA} \, ,} \tag{17}$$

where $\theta(R-r)$ is the Heaviside function taking the unit value for $r < R$ and equal to zero for $r > R$. We note that the solution should be continuous with its derivative at $r = R$: $c_{SA}(R-0,t) = c_{SA}(R+0,t)$ and $\partial_r c_{SA}(R-0,t) = \partial_r c_{SA}(R+0,t)$.

Eq. 17 can also be interpreted as an adiabatic approximation of our starting system of PDEs (Eq. 1). Specifically, by assuming that at a given concentration of SA, $c_{SA}$, the density of bridges instantaneously relaxes towards its equilibrium value, we find

$$K_{on}\left(1-\frac{c_B}{c_{HA}^0}\right)c_{SA} = k_{off}\,c_B\,. \tag{18}$$

By using the previous equation in the first of Eq. 1, we recover Eq. 17.

An important remark is that we can rescale the spatial and temporal coordinates according to

$$\mathbf{r} \equiv R\,\mathbf{r}_* \qquad \text{and} \qquad t \equiv T\,t_* \qquad \text{with} \qquad T \equiv \frac{R^2}{D} \tag{19}$$

into new dimensionless coordinates $\mathbf{r}_*$ and $t_*$, so that the equation of motion becomes

$$\partial_{t_*} c_{SA} = \nabla_*^2 c_{SA} - \lambda\,\theta(1-r_*)\,c_{SA} \qquad \text{with} \qquad \lambda \equiv \frac{K_d R^2}{D} \tag{20}$$

and the Neumann boundary condition $\partial_{r_*} c_{SA}(r_* = \ell, t_*) = 0$ at $\ell \equiv L/R$. Therefore, the mathematical solution of the problem only depends on the two dimensionless parameters

$$\lambda \equiv \frac{K_d R^2}{D} \qquad \text{and} \qquad \ell \equiv \frac{L}{R} > 1\,. \tag{21}$$

#### 2.3 The short–time decay

The early decay is controlled by the rate $K_d$, giving

$$c_{SA}(r,t) \simeq c_{SA}^0\, e^{-K_d t} \tag{22}$$

as observed in the numerical computations (Main Fig. 4b). Discrepancies are because SA replenishment is already at play during the early stage of the dynamics, as shown in Fig. S2. Over longer time scales, the decay is controlled by diffusion, for which Eq. 17 should be solved.

#### 2.4 Solving the partial differential equation for $c_{SA}$ (Eq. 17) in 2D

In 2D, the Laplacian is given by

$$\nabla^2 c = \partial_r^2 c + r^{-1}\,\partial_r c + r^{-2}\,\partial_\theta^2 c\,. \tag{23}$$

Since the partial differential equation is (piecewise) linear, its general solution can be decomposed in terms of eigenfunctions and their associated eigenvalues. Since the initial conditions and the boundary conditions have the circular symmetry, the eigenfunctions to consider should not depend on the angle $\theta$ and should obey the following eigenvalue equation:

$$\hat{L}\phi \equiv D\left(\frac{d^2\phi}{dr^2} + \frac{1}{r}\frac{d\phi}{dr}\right) - K_d\,\theta(R-r)\,\phi = -\gamma\,\phi\,. \tag{24}$$

Furthermore, we note that the linear operator is Hermitean: $\hat{L} = \hat{L}^\dagger$. Therefore, the eigenvalues are real and the eigenfunctions can be used at left and at right of the expansion. Moreover, the eigenfunctions can be taken as real functions.

The general solution can thus be written as

$$c_{SA}(r,t) = \sum_n \mathcal{C}_n\, e^{-\gamma_n t}\,\phi_n(r) \qquad \text{with} \qquad \mathcal{C}_n = \frac{\int c_{SA}(r,0)\,\phi_n(r)\,d^2r}{\int [\phi_n(r)]^2\,d^2r}\,, \tag{25}$$

where the spatial integration extends to the domain $r < L$.

The eigenvalue equation reads

$$\frac{d^2\phi_n}{dr^2} + \frac{1}{r}\frac{d\phi_n}{dr} = -\frac{\gamma_n - K_d}{D}\,\phi_n \qquad (r < R)\,, \tag{26}$$
$$\frac{d^2\phi_n}{dr^2} + \frac{1}{r}\frac{d\phi_n}{dr} = -\frac{\gamma_n}{D}\,\phi_n \qquad (r > R)\,, \tag{27}$$

with the boundary conditions

$$\phi_n(R-0) = \phi_n(R+0)\,, \qquad \frac{d\phi_n}{dr}(R-0) = \frac{d\phi_n}{dr}(R+0)\,, \qquad \text{and} \qquad \frac{d\phi_n}{dr}(L) = 0\,. \tag{28}$$

The solutions are given by Bessel functions or modified Bessel functions of zeroth order.

For $\gamma < K_d$, the eigenvalue is given by

$$\gamma = D\,q^2 \tag{29}$$

and the eigenfunction by

$$\phi(r) = C\,I_0(\kappa r) \qquad (r < R)\,, \tag{30}$$

$$\phi(r) = A\,J_0(qr) + B\,Y_0(qr) \qquad (r > R)\,, \tag{31}$$

with

$$\kappa = \sqrt{\frac{K_d}{D} - q^2}\,. \tag{32}$$

The boundary conditions at $r = R$ imply that

$$\begin{pmatrix} A \\ B \end{pmatrix} = \frac{C}{W}\begin{pmatrix} Y_0'(qR) & -Y_0(qR) \\ -J_0'(qR) & J_0(qR) \end{pmatrix}\begin{pmatrix} I_0(\kappa R) \\ \frac{\kappa}{q}\,I_0'(\kappa R) \end{pmatrix} \tag{33}$$

with the Wronskian

$$W = J_0(qR)\, Y_0'(qR) - Y_0(qR)\, J_0'(qR) = \frac{2}{\pi q R}\,. \tag{34}$$

For $\gamma > K_d$, the eigenvalue is again given by

$$\gamma = D\, q^2 \tag{35}$$

but the eigenfunction by

$$\phi(r) = C\, J_0(kr) \qquad\qquad (r < R)\,, \tag{36}$$

$$\phi(r) = A\, J_0(qr) + B\, Y_0(qr) \qquad (r > R)\,, \tag{37}$$

with

$$k = \sqrt{q^2 - \frac{K_d}{D}}\,. \tag{38}$$

The boundary conditions at $r = R$ imply that

$$\begin{pmatrix} A \\ B \end{pmatrix} = \frac{C}{W} \begin{pmatrix} Y_0'(qR) & -Y_0(qR) \\ -J_0'(qR) & J_0(qR) \end{pmatrix} \begin{pmatrix} J_0(kR) \\ \frac{k}{q}\, J_0'(kR) \end{pmatrix} \tag{39}$$

with the same Wronskian as above.

In both cases, the eigenvalues are obtained from the Neumann boundary condition

$$A\, J_0'(q_n L) + B\, Y_0'(q_n L) = 0 \tag{40}$$

with the respective coefficients $A$ and $B$.

#### 2.5 The long-time limit

The long-time limit is controlled by the smallest wavenumber $q$. Therefore, this limit is controlled by eigenvalues with $\gamma \ll K_d$, $q \ll \sqrt{K_d/D}$, and such that $\kappa \simeq \sqrt{K_d/D}$. On the longest time scale, the decay is exponential and given by

$$c_{SA}(r,t) \simeq \mathcal{C}\, e^{-\gamma t}\, \phi(r) \qquad \text{with} \qquad \mathcal{C} = \frac{\int c_{SA}(r,0)\, \phi(r)\, d^2r}{\int [\phi(r)]^2\, d^2r}\,, \tag{41}$$

where $\gamma = Dq^2$ is the smallest non-zero eigenvalue and $\phi(r)$ the corresponding eigenfunction. This eigenvalue is given by the smallest non-zero root of

$$A\, J_0'(qL) + B\, Y_0'(qL) = 0\,. \tag{42}$$

In general, the coefficients $A$ and $B$ depend on all the parameters of the problem, including $R$, $K_d$, and $D$, so that the roots $q_n$ also depend on these parameters. For the same reason, the eigenfunction will depend on the diffusion coefficient and is complicated to obtain. We note that, in the problem at hand, we have $\frac{K_d}{D} R^2 \simeq O(1)$, so that the results will depend on all the parameters in a complicated way.

The integral

$$N_{SA}(t) \equiv \int_{r<R} c_{SA}(r,t)\, d^2r \tag{43}$$

should give the the number of bridges according to

$$N_B(t) \simeq \frac{K_{on}}{k_{off}}\, N_{SA}(t)\,. \tag{44}$$

In the long-time limit, the exponential decay goes as

$$N_{SA}(t) \propto e^{-Dq^2 t}\,, \tag{45}$$

so that

$$N_B(t) \simeq \frac{K_{on}}{k_{off}} \, N_{SA}(t) \propto \frac{K_{on}}{k_{off}} \, e^{-Dq^2 t} \tag{46}$$

for $t \to \infty$ in a finite system.

The issue is to evaluate $q$ and $\gamma$ in terms of the parameters of the problem, namely, $D$, $K_d$, $R$, and $L$. This problem is addressed in the next two sections.

#### 2.6 The limit $\lambda = K_d R^2/D \ll 1$

In this limit, we can use the approximations of the Bessel functions for $z \ll 1$:

$$I_0(z) = 1 + \frac{z^2}{4} + O(z^4)\,, \tag{47}$$

$$J_0(z) = 1 - \frac{z^2}{4} + O(z^4)\,, \tag{48}$$

$$Y_0(z) = \frac{2}{\pi}\left(\ln\frac{z}{2} + \gamma\right) - \frac{z^2}{2\pi}\left(\ln\frac{z}{2} + \gamma - 1\right) + O(z^4 \ln z)\,, \tag{49}$$

so that

$$I_0'(z) = \frac{z}{2} + O(z^3)\,, \tag{50}$$

$$J_0'(z) = -\frac{z}{2} + O(z^3)\,, \tag{51}$$

$$Y_0'(z) = \frac{2}{\pi z} - \frac{z}{\pi}\left(\ln\frac{z}{2} + \gamma - \frac{1}{2}\right) + O(z^3 \ln z)\,. \tag{52}$$

As a consequence, the coefficients $A$ and $B$ can be expanded to give

$$A = C\,(1 + \alpha\,\lambda + \cdots) \qquad \text{with} \qquad \alpha = -\frac{1}{2}\left(\ln\frac{qR}{2} + \gamma - \frac{1}{2}\right)\,, \tag{53}$$

$$B = C\,(\beta\,\lambda + \cdots) \qquad \text{with} \qquad \beta = \frac{\pi}{4}\,. \tag{54}$$

Therefore, the Neumann boundary condition

$$A\,J_0'(qL) + B\,Y_0'(qL) = 0 \tag{55}$$

becomes

$$-(qL)^2 + \lambda + \frac{\lambda}{2}\,(qL)^2\,\ln\frac{L}{R} \simeq 0\,. \tag{56}$$

Since the size $L$ of the domain is always larger than the radius $R$ of the contact zone of the virus, we may consider the limit $L \gg R$, so that

$$q \simeq \frac{1}{L}\sqrt{\frac{2}{\ln\frac{L}{R}}} \qquad \text{for} \qquad L \gg R\,. \tag{57}$$

Finally, the leading eigenvalue of the problem with the Neumann boundary condition is thus given by

$$\boxed{\gamma = Dq^2 \simeq \frac{2\,D}{L^2\ln\frac{L}{R}} \qquad \text{for} \qquad \lambda = \frac{K_d R^2}{D} \ll 1 \qquad \text{and} \qquad L \gg R\,.} \tag{58}$$

#### 2.7 The limit $L \gg R$ only

For arbitrary values of $\lambda = K_d R^2/D$, we have

$$A \simeq C\left[I_0(\sqrt{\lambda}) - \sqrt{\lambda}\,I_0'(\sqrt{\lambda})\left(\ln\frac{qR}{2} + \gamma\right)\right], \tag{59}$$

$$B \simeq C\,\frac{\pi}{2}\,\sqrt{\lambda}\,I_0'(\sqrt{\lambda})\,. \tag{60}$$

In the limit $qR \ll 1$, we get

$$A \simeq B\,\frac{2}{\pi}\left(\ln\frac{qR}{2} + \gamma\right). \tag{61}$$

Substituting this result into the Neumann boundary condition

$$A\,J_0'(qL) + B\,Y_0'(qL) = 0 \tag{62}$$

and using the formula

$$Y_0(z) = \frac{2}{\pi}\,J_0(z)\left(\ln\frac{z}{2} + \gamma\right) + \frac{z^2}{2\pi} + O(z^4)\,, \tag{63}$$

for $z \ll 1$, we find

$$2\,J_0'(qL)\,\ln\frac{L}{R}+\frac{2}{qL}\,J_0(qL)+qL\simeq 0\,. \tag{64}$$

Since $J_0(z)=1-z^2/4+O(z^4)$ and $J_0'(z)=-z/2+O(z^3)$, we again obtain

$$q\simeq\frac{1}{L}\sqrt{\frac{2}{\ln\frac{L}{R}}}\qquad\text{for}\qquad L\gg R\,. \tag{65}$$

Therefore, we can confirm that the leading eigenvalue of the problem with the Neumann boundary condition is given by

$$\boxed{\gamma=Dq^2\simeq\frac{2\,D}{L^2\ln\frac{L}{R}}\qquad\text{for}\qquad\lambda=\frac{K_dR^2}{D}\ll 1\qquad\text{and}\qquad L\gg R\,.} \tag{66}$$

### 3 Asymptotic analysis of the mean-field theory: Dirichlet boundary conditions

We aim to solve Eqs. 1 with boundary condition

$$c_{SA}(r=L,t)=c^0_{SA} \tag{67}$$

at the border $r=L$ of the domain of interest. The parameters are defined as in Eqs. 5. The initial conditions and the peak maximum are estimated as in Sec. 2.1.

#### 3.1 The stationary solution

For Dirichlet boundary conditions, the stationary solution is not identically equal to zero. Instead, the stationary solution should obey

$$\partial_t\,c_B=K_{on}\left(1-\frac{c_B}{c^0_{HA}}\right)c_{SA}-k_{\mathit{off}}\,c_B=0 \tag{68}$$

for $r > R$. Substituting this result into the equation for $\partial_t c_{SA} = 0$, we find

$$D\nabla^2 c_{SA} - K_d\, c_{SA} = 0 \tag{69}$$

for $r < R$, and

$$D\nabla^2 c_{SA} = 0 \tag{70}$$

for $R < r < L$. At $r = R$, the function $c_{SA}$ should be continuous with its derivative $\partial_r c_{SA}$.

Therefore, the stationary solution of this problem is given in terms of the modified Bessel function of zeroth order $I_0(z)$ by

$$c_{SA}(r) = A\, I_0\!\left(r\sqrt{\frac{K_d}{D}}\right) \qquad \text{for} \quad r < R\,, \tag{71}$$

$$c_{SA}(r) = c_{SA}^0 - B\, \ln\frac{L}{r} \qquad \text{for} \quad R < r < L\,, \tag{72}$$

with the coefficients

$$A = \frac{c_{SA}^0}{I_0\!\left(R\sqrt{\frac{K_d}{D}}\right) + I_0'\!\left(R\sqrt{\frac{K_d}{D}}\right) R\,\sqrt{\frac{K_d}{D}}\, \ln\frac{L}{R}}\,, \tag{73}$$

$$B = A\, I_0'\!\left(R\sqrt{\frac{K_d}{D}}\right) R\,\sqrt{\frac{K_d}{D}}\,, \tag{74}$$

and

$$c_B = \frac{K_{on}\, c_{SA}}{k_{off} + K_{on}\dfrac{c_{SA}}{c_{HA}^0}}\,. \tag{75}$$

The stationary profile for $c_{SA}(r)$ agrees with the numerical observations, since $I_0(z) \simeq 1 + z^2/4$ for small values of $z$.

#### 3.2 Linearization around the stationary solution

In order to understand the long-time dynamics, the equations should be linearized around the stationary solution according to $c_{SA} = c_{SA,st} + \delta c_{SA}$. The problem is complicated by the spatial dependence of the stationary solution. If we may assume that

$$k_{off} \gg K_{on} \, \frac{c^0_{SA}}{c^0_{HA}} \, , \tag{76}$$

we can simplify the problem because

$$c_B \simeq \frac{K_{on}}{k_{off}} \, c_{SA} \, , \tag{77}$$

so that the linear equations to be solved reduce to

$$\partial_t \, \delta c_{SA} = D \nabla^2 \delta c_{SA} - K_d \, \theta(R - r) \, \delta c_{SA} \, , \tag{78}$$

where $\theta(R - r)$ is the Heaviside function taking the unit value for $r < R$ and equal to zero for $r > R$, and

$$\delta c_B \simeq \frac{K_{on}}{k_{off}} \, \delta c_{SA} \, . \tag{79}$$

An important remark is that we can rescale the spatial and temporal coordinates according to

$$\mathbf{r} \equiv R \, \mathbf{r}_* \qquad \text{and} \qquad t \equiv T \, t_* \qquad \text{with} \qquad T \equiv \frac{R^2}{D} \tag{80}$$

into new dimensionless coordinates $\mathbf{r}_*$ and $t_*$, so that the equation of motion becomes

$$\partial_{t_*} \, \delta c_{SA} = \nabla^2_* \delta c_{SA} - \lambda \, \theta(1 - r_*) \, \delta c_{SA} \qquad \text{with} \qquad \lambda \equiv \frac{K_d R^2}{D} \tag{81}$$

and the Dirichlet boundary condition $\delta c_{SA}(r_* = \ell, t_*) = 0$ at $\ell \equiv L/R$. Therefore, the mathematical solution of this linear problem only depends on the two dimensionless parameters

$$\lambda \equiv \frac{K_d R^2}{D} \qquad \text{and} \qquad \ell \equiv \frac{L}{R} > 1\,. \tag{82}$$

We note that the early decay is controlled by the decay rate $K_d$, giving

$$\delta c_{SA}(r,t) \simeq \delta c_{SA}^0 \, e^{-K_d t} \tag{83}$$

as observed in the numerical computations (see Sec. 2.3). Over longer time scales, the decay is controlled by diffusion, for which the partial differential equation should be solved.

##### 3.3 Solving the partial differential equation for $\delta c_{SA}$ in 2D

As done in Sec. 2.4, we need to solve the following eigenvalue equation:

$$\hat{L}\phi \equiv D\left(\frac{d^2\phi}{dr^2} + \frac{1}{r}\frac{d\phi}{dr}\right) - K_d\,\theta(R-r)\,\phi = -\gamma\,\phi\,. \tag{84}$$

The general solution can be written as

$$\delta c_{SA}(r,t) = \sum_n \mathcal{C}_n\, e^{-\gamma_n t}\,\phi_n(r) \qquad \text{with} \qquad \mathcal{C}_n = \frac{\int c_{SA}(r,0)\,\phi_n(r)\,d^2r}{\int [\phi_n(r)]^2\,d^2r}\,, \tag{85}$$

where the spatial integration extends to the domain $r < L$.

The eigenvalue equation reads

$$\frac{d^2\phi_n}{dr^2} + \frac{1}{r}\frac{d\phi_n}{dr} = -\frac{\gamma_n - K_d}{D}\,\phi_n \qquad (r < R)\,, \tag{86}$$

$$\frac{d^2\phi_n}{dr^2} + \frac{1}{r}\frac{d\phi_n}{dr} = -\frac{\gamma_n}{D}\,\phi_n \qquad (r > R)\,, \tag{87}$$

with the boundary conditions

$$\phi_n(R-0) = \phi_n(R+0)\,, \qquad \frac{d\phi_n}{dr}(R-0) = \frac{d\phi_n}{dr}(R+0)\,, \qquad \text{and} \qquad \phi_n(L) = 0\,. \tag{88}$$

The solutions are given by Bessel functions or modified Bessel functions of zeroth order.

For $\gamma < K_d$, the eigenvalue is given by

$$\gamma = D\,q^2 \tag{89}$$

and the eigenfunction by

$$\phi(r) = C\,I_0(\kappa r) \qquad (r < R)\,, \tag{90}$$

$$\phi(r) = A\,J_0(qr) + B\,Y_0(qr) \qquad (r > R)\,, \tag{91}$$

with

$$\kappa = \sqrt{\frac{K_d}{D} - q^2}\,. \tag{92}$$

The boundary conditions at $r = R$ imply that

$$\begin{pmatrix} A \\ B \end{pmatrix} = \frac{C}{W} \begin{pmatrix} Y_0'(qR) & -Y_0(qR) \\ -J_0'(qR) & J_0(qR) \end{pmatrix} \begin{pmatrix} I_0(\kappa R) \\ \frac{\kappa}{q}\,I_0'(\kappa R) \end{pmatrix} \tag{93}$$

with the Wronskian

$$W = J_0(qR)\,Y_0'(qR) - Y_0(qR)\,J_0'(qR) = \frac{2}{\pi q R}\,. \tag{94}$$

For $\gamma > K_d$, the eigenvalue is again given by

$$\gamma = D\,q^2 \tag{95}$$

but the eigenfunction by

$$\phi(r) = C\,J_0(kr) \qquad\qquad (r < R)\,, \tag{96}$$

$$\phi(r) = A\,J_0(qr) + B\,Y_0(qr) \qquad (r > R)\,, \tag{97}$$

with

$$k = \sqrt{q^2 - \frac{K_d}{D}}\,. \tag{98}$$

The boundary conditions at $r = R$ imply that

$$\begin{pmatrix} A \\ B \end{pmatrix} = \frac{C}{W} \begin{pmatrix} Y_0'(qR) & -Y_0(qR) \\ -J_0'(qR) & J_0(qR) \end{pmatrix} \begin{pmatrix} J_0(kR) \\ \frac{k}{q}\,J_0'(kR) \end{pmatrix} \tag{99}$$

with the same Wronskian as above.

In both cases, the eigenvalues are obtained from the Dirichlet boundary condition

$$A\,J_0(q_n L) + B\,Y_0(q_n L) = 0 \tag{100}$$

with the respective coefficients $A$ and $B$.

#### 3.4 The long-time limit

The longer the time $t$, the smaller the wavenumber $q$. Therefore, the long-time behavior is controlled by eigenvalues with $\gamma \ll K_d$, $q \ll \sqrt{K_d/D}$, and such that $\kappa \simeq \sqrt{K_d/D}$. On the longest time scale, the decay is exponential and given by

$$\delta c_{SA}(r,t) \simeq \mathcal{C}\,e^{-\gamma t}\,\phi(r) \qquad \text{with} \qquad \mathcal{C} = \frac{\int c_{SA}(r,0)\,\phi(r)\,d^2r}{\int [\phi(r)]^2\,d^2r}\,, \tag{101}$$

where $\gamma = Dq^2$ is the smallest non-zero eigenvalue and $\phi(r)$ the corresponding eigenfunction. This eigenvalue is given by the smallest non-zero root of

$$A\, J_0(qL) + B\, Y_0(qL) = 0\,. \tag{102}$$

In general, the coefficients $A$ and $B$ depend on all the parameters of the problem, including $R$, $K_d$, and $D$, so that the roots $q_n$ also depend on these parameters. For the same reason, the eigenfunction will depend on the diffusion coefficient and is complicated to obtain. We note that, in the problem at hand, we have $\frac{K_d}{D} R^2 \simeq O(1)$, so that the results will depend on all the parameters in a complicated way.

The integral

$$N_{SA}(t) \equiv \int_{r<R} c_{SA}(r,t)\, d^2 r \tag{103}$$

should give the number of bridges according to

$$N_B(t) \simeq \frac{K_{on}}{k_{off}}\, N_{SA}(t)\,. \tag{104}$$

Since $c_{SA} = c_{SA,st} + \delta c_{SA}$, we find that

$$N_{SA}(t) = N_{SA,st} + \mathcal{N}\, e^{-Dq^2 t}\,, \tag{105}$$

so that

$$N_B(t) \simeq \frac{K_{on}}{k_{off}}\, N_{SA}(t) \simeq \frac{K_{on}}{k_{off}} \left( N_{SA,st} + \mathcal{N}\, e^{-Dq^2 t} \right) \tag{106}$$

in the long-time limit $t \to \infty$ for a finite system.

The issue is to evaluate $q$ and $\gamma$ in terms of the parameters of the problem, namely, $D$, $K_d$, $R$, and $L$.

##### 3.5 The limit $L \gg R$

Using the approximations of the Bessel functions for $z = qR \ll 1$:

$$J_0(z) = 1 - \frac{z^2}{4} + O(z^4)\,, \tag{107}$$

$$Y_0(z) = \frac{2}{\pi}\left(\ln\frac{z}{2} + \gamma\right) - \frac{z^2}{2\pi}\left(\ln\frac{z}{2} + \gamma - 1\right) + O(z^4 \ln z)\,, \tag{108}$$

so that

$$J_0'(z) = -\frac{z}{2} + O(z^3)\,, \tag{109}$$

$$Y_0'(z) = \frac{2}{\pi z} - \frac{z}{\pi}\left(\ln\frac{z}{2} + \gamma - \frac{1}{2}\right) + O(z^3 \ln z)\,, \tag{110}$$

but for arbitrary values of $\lambda = K_d R^2/D$, we have

$$A \simeq C\left[I_0(\sqrt{\lambda}) - \sqrt{\lambda}\, I_0'(\sqrt{\lambda})\left(\ln\frac{qR}{2} + \gamma\right)\right], \tag{111}$$

$$B \simeq C\,\frac{\pi}{2}\sqrt{\lambda}\, I_0'(\sqrt{\lambda})\,, \tag{112}$$

so that

$$A \simeq B\,\frac{2}{\pi}\left(\ln\frac{qR}{2} + \gamma\right). \tag{113}$$

Substituting this result into the Dirichlet boundary condition

$$A\, J_0(qL) + B\, Y_0(qL) = 0 \tag{114}$$

and using the formula

$$Y_0(z) = \frac{2}{\pi}\, J_0(z)\left(\ln\frac{z}{2} + \gamma\right) + \frac{z^2}{2\pi} + O(z^4)\,, \tag{115}$$

for $z \ll 1$, we find

$$J_0(qL) \simeq -\frac{(qL)^2}{4\,\ln\frac{L}{R}}\,. \tag{116}$$

Since $\ln(L/R) \to \infty$ for $L/R \to \infty$, the root of this equation corresponds in this limit to the root of $J_0(qL) \simeq 0$, which is nothing but the first root of the Bessel function $J_0(z)$, i.e., $z = j_{0,1} \simeq 2.40483$. Therefore, we obtain

$$q \simeq \frac{j_{0,1}}{L} \simeq \frac{2.40483}{L}\,. \tag{117}$$

so that the leading eigenvalue of the problem with the Dirichlet boundary condition is given by

$$\boxed{\gamma = Dq^2 \simeq D\left(\frac{2.40483}{L}\right)^2 \qquad \text{for} \qquad L \gg R\,,} \tag{118}$$

as it should.

### 4 Distribution of the detachment time, $P_{detach}(t_{detach})$

We define by $p_i$ the probability of having $i$ bridges ($n_b = i$). The evolution of $p_i$ is given by the following master equation

$$\frac{dp_i}{dt} \;=\; g_{i-1}(t)p_{i-1} + r_{i+1}p_{i+1} - (g_i(t) + r_i)p_i\,, \tag{119}$$

where $r_i$ and $g_i$ are the transition rates towards a state with, respectively, $i-1$ and $i+1$ bridges.[2,3] We employ a mean field approach in which we average over the concentration of SA underneath the disc. The rates are then given by

$$r_i \;=\; k_{off} \cdot i \tag{120}$$

$$g_i(t) \;=\; k_{on}\pi\lambda^2(N_{HA} - i)\langle c_{SA}\rangle(t)\,, \tag{121}$$

where

$$\langle c_{SA}\rangle(t) \;=\; \frac{2}{R^2}\int_0^R dr\, r\cdot c_{SA}(r,t)\,. \tag{122}$$

Using mean first-passage theory,[2] we can calculate the average time taken by the system to reach the $n_b = 0$ state starting from $n_b = m$ at $t = 0$:

$$T_{m,0}(t) \;=\; \sum_{i=1}^{m}\left[\frac{1}{r_i(t)} + \sum_{j=i+1}^{N_{HA}}\frac{1}{r_j(t)}\prod_{k=1}^{j-1}\frac{g_k(t)}{r_k(t)}\right] \tag{123}$$

The average time to detach at a given $c_{SA}$, $\tau_d$ ($\tau_d = 1/k_{detach}$, see Main Eq. 4), is then calculated by averaging $T_{m,0}$ using the equilibrium probability of having $m$ bridges ($p_m$)

$$k_{det}(t) \;=\; \left[\sum_{m=1}^{N_{NA}} p_m(t)T_{m,0}(t)\right]^{-1}\,. \tag{124}$$

$p_m$ can be calculated recursively using

$$p_m = p_{m-1}\frac{g_{m-1}}{r_m}\,. \tag{125}$$

### 5 Supplementary figures

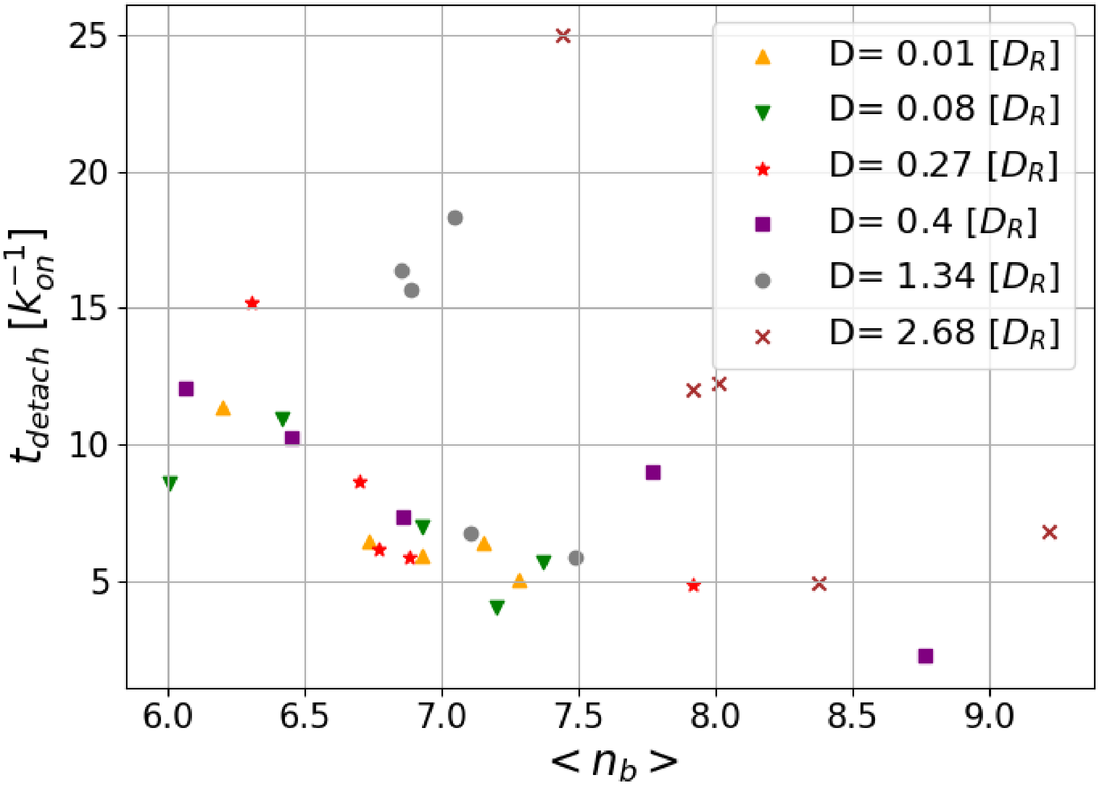


Figure S3: Detachment times sampled by simulations in which the particle diffuses with a constant diffusion constant $D$ (instead of $D = D_R/n_b$). Simulation parameters: $k_d = 3.571\, k_{on}$, $k_{off} = 16.07\, k_{on}$, $D_R = 0.3\, \mu\mathrm{m}^2$.

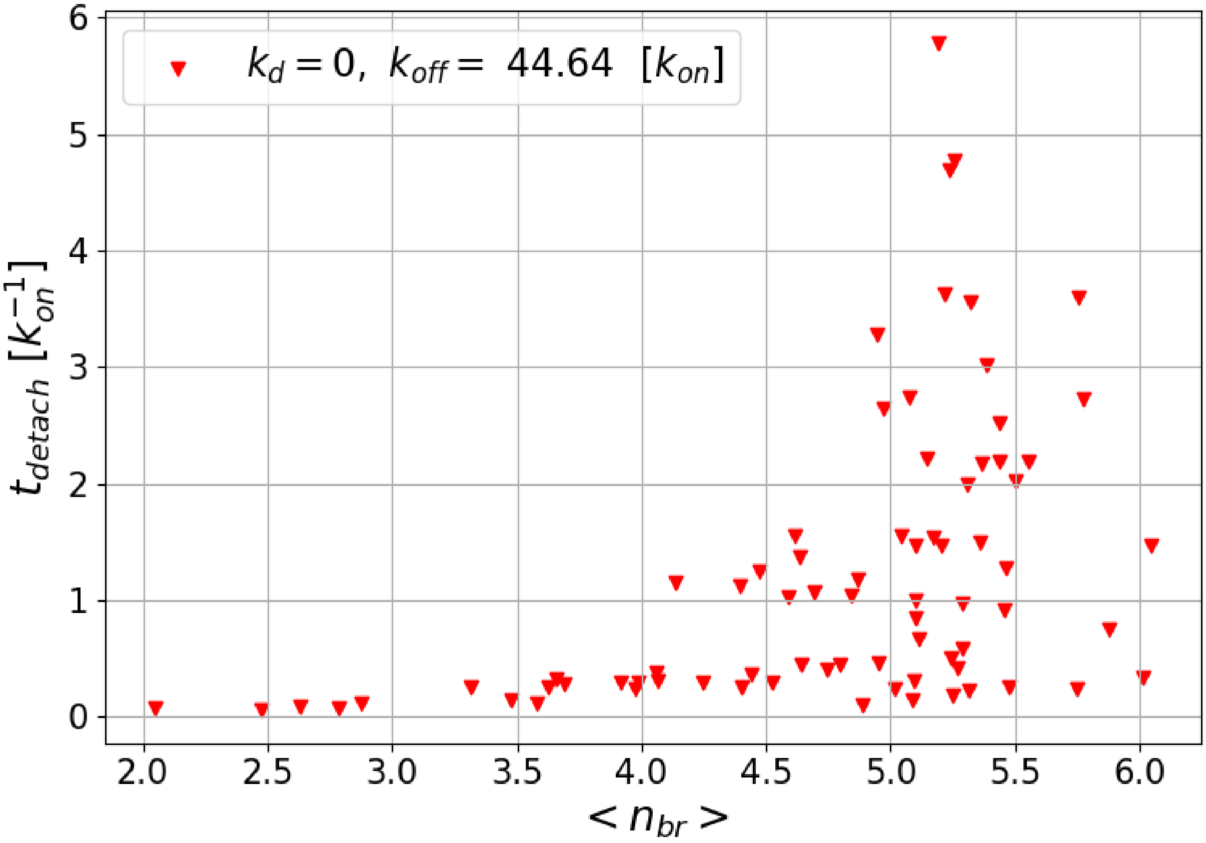


Figure S4: Detachment time without NA activity. As compared to Main Fig. 2, we increased the value of $k_{off}$ to keep the simulations affordable.

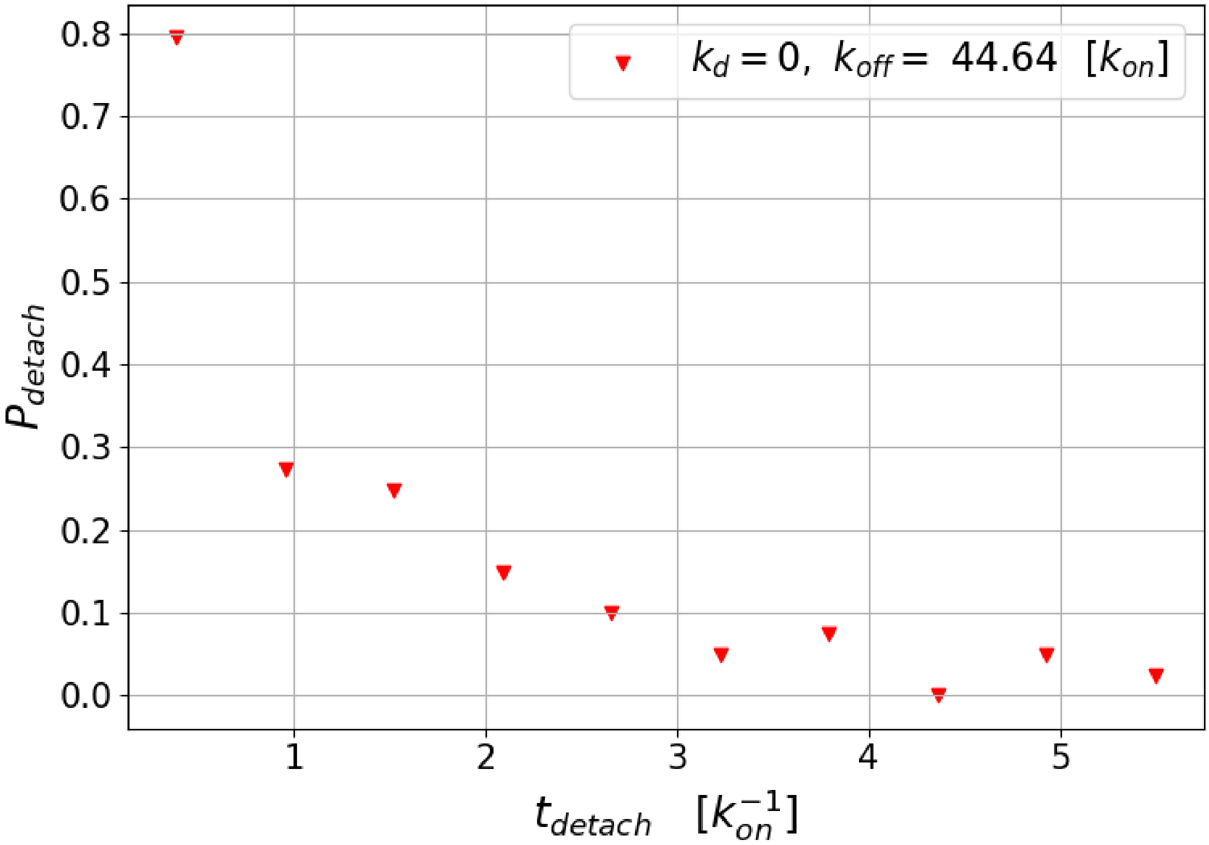


Figure S5: Distribution of the detachment times reported in Fig. S4.

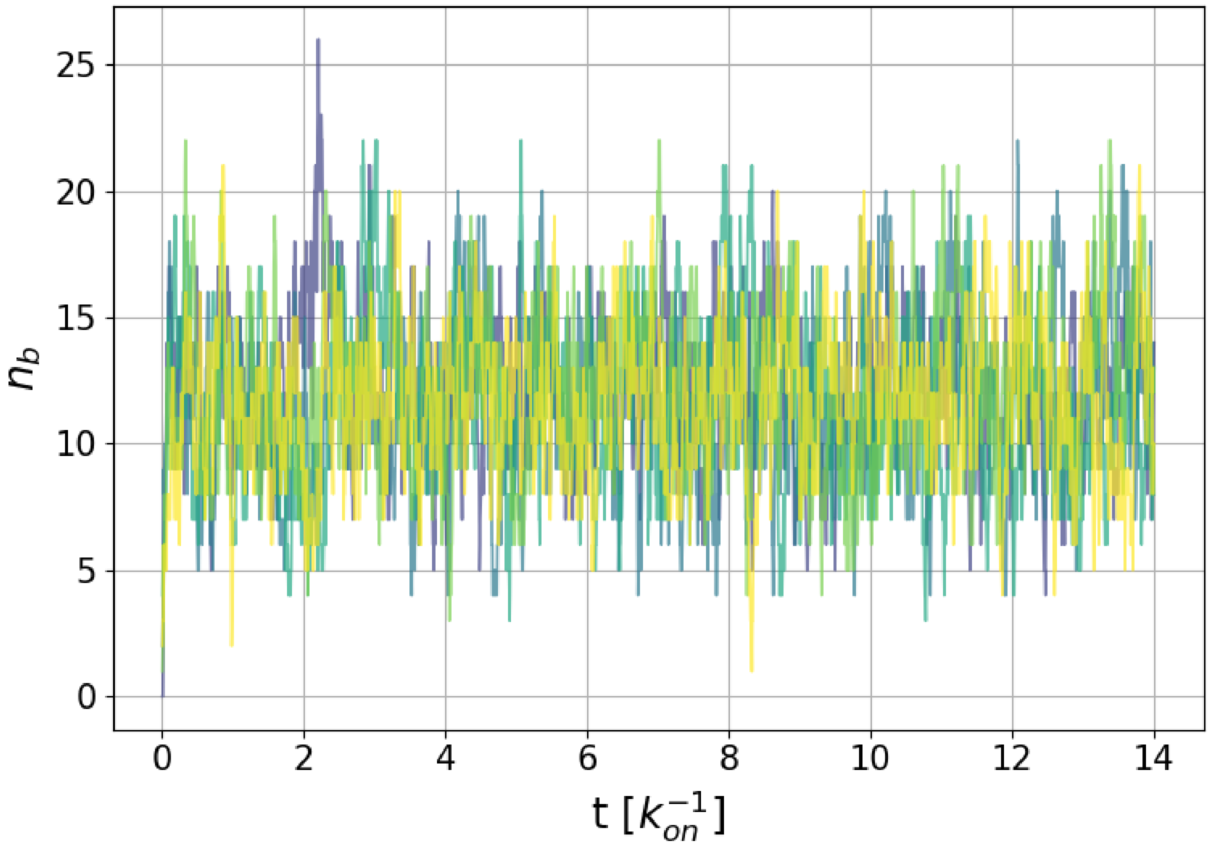


Figure S6: Number of bridges *vs* time for $k_{off} = 16.07\, k_{on}$ and $k_d = 0$.

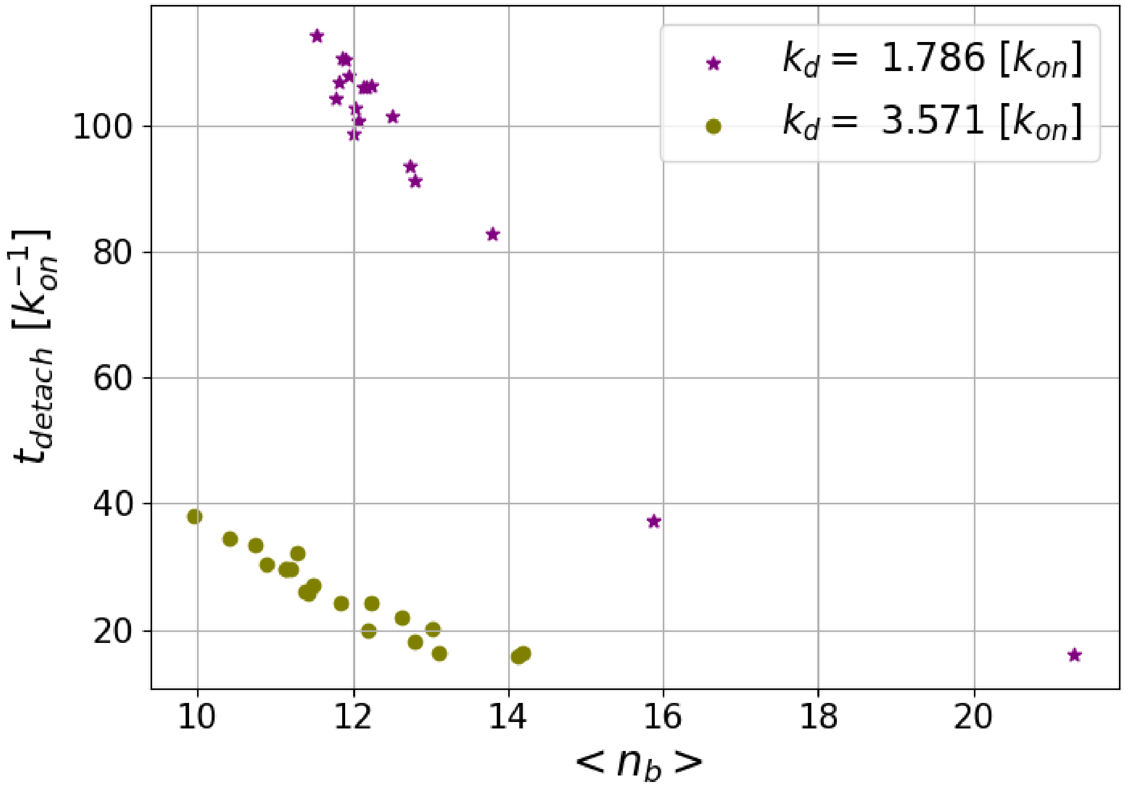


Figure S7: Detachment times for a particle with $R = 120\,\text{nm}$ ($N_{NA} = 51$ and $N_{HA} = 190$) and $k_{off} = 16.07\,k_{on}$.

### References


(1) Muller, M.; Lauster, D.; Wildenauer, H. H.; Herrmann, A.; Block, S. Mobility-based quantification of multivalent virus-receptor interactions: new insights into influenza A virus binding mode. *Nano letters* **2019**, *19*, 1875–1882.

(2) Van kampen, N. G. In *Stochastic Processes in Physics and Chemistry (Third Edition)*; Van kampen, N. G., Ed.; North-Holland Personal Library; Elsevier, pp 292–325.

(3) Erdmann, T.; Schwarz, U. S. Impact of receptor-ligand distance on adhesion cluster stability. *22*, 123–137.
